# A framework for comparing microbial networks reveals core associations

**DOI:** 10.1101/2020.10.05.325860

**Authors:** Lisa Röttjers, Doris Vandeputte, Jeroen Raes, Karoline Faust

## Abstract

Microbial network construction and analysis is an important tool in microbial ecology. As microbial interactions are challenging to infer experimentally, such networks are often constructed from statistically inferred associations and may not represent ecological interactions. Hence, microbial association networks contain a large number of errors and their derived properties do not necessarily reflect true community structure. Such errors can be identified with the use of appropriate null models. We have developed anuran, a toolbox for investigation of noisy networks with null models, for identification of non-random patterns in groups of association networks. This toolbox compares multiple networks to identify conserved subsets (core association networks, CANs) and other network properties that are shared across all networks. Such groups of networks can be generated from a collection of time series data or from cross-sectional sample sets. We use data from the Global Sponge Project to demonstrate that different orders of sponges have a larger CAN than expected at random.

## Introduction

A biologically interesting pattern needs to differ from patterns observed by chance, or from patterns generated by processes that are not of interest to the investigator (Strong, 1980). Such differences can often only be observed by generating data under the sets of rules specified by null models. For network inference, tools such as CoNet and LSA use a conceptually simple null model where shuffled data is presumed to represent the situation without meaningful biological structure (Faust & Raes, 2016; Ruan et al., 2006). In the analysis of microbial networks however, null models are not yet systematically employed, so that network properties are often misinterpreted as being due to a process of interest instead of resulting from properties of the count table or for other reasons. Here, we present a tool for generation of randomized networks through null models that respect important characteristics of the original network.

Through null models, researchers can assess whether derived network properties could be observed even if the network is a random collection of edges, rather than biologically meaningful taxon-taxon associations. If network properties such as hub nodes, cluster coefficients or degree distributions are meaningful, they should be different from their estimates for randomized data. Since microbial networks are known to be inaccurate (Hirano & Takemoto, 2019; Röttjers & Faust, 2018; Weiss et al., 2016), there is a risk that network properties may originate from other processes rather than the biological process(es) of interest. Null models can help identify network properties that are different from what is expected based on the null hypothesis.

One of those properties of interest is the edge intersection of networks, the group of edges present in a combination of networks. Comparisons across multiple networks have previously been used to identify meaningful associations. For example, Wang et al. computed the similarities between networks inferred with different methods to identify meaningful associations (Wang et al., 2017). Similarly, Jackson et al. compared networks from different populations and used null models to show that gut microbial association networks from geographically separated populations were more similar than expected by chance (Jackson et al., 2018). We expect more network comparison studies in the future; such studies could include different populations, treatment effects, or separate time series.

For these comparisons in particular, null models are crucial to identify similarities across networks not driven by similarity in species composition. The edge intersection of networks can become large even for completely random networks due to similarities in the abundance data. These similarities are relevant when ecosystems consist of a number of “core” genera that are found in most samples of the ecosystem. Such core microbiomes have been identified for the human gut (Falony et al., 2016), the oral microbiome (Zaura et al., 2009) and the coral microbiome (Ainsworth et al., 2015). Yet, presence of a core microbiome does not necessarily imply presence of a core microbial interaction network, since the same microorganisms may interact in different ways, fluctuations of their abundances may not be driven by interactions, or the data may be too noisy to infer associations. This raises the question whether interactions are preserved across different realisations of an ecosystem, whether they are preserved within sub-groups of the ecosystem or whether they are unique (Bashan et al., 2016). Since null model analysis can distinguish random from significant intersection sizes, it can test for the presence of one or more core association networks (CANs) within a group of association networks. Core associations can then be further explored to check whether they represent ecological interactions. In this way, null model analysis is a step towards answering whether interactions are universal.

For investigation of non-random network properties, several network null models have already been implemented in previous research or in software. Prior work demonstrated that many network properties can be highly correlated, but may represent different underlying aspects in network topology. For example, Valente et al. (2008) found strong correlations between degree and several other centralities, including eigenvector and betweenness centrality in their analysis of 62 sociometric networks (Valente et al., 2008). For bipartite networks, Dormann et al. (2009) demonstrated that the degree and edge density were strongly correlated to species number, while the cluster coefficient and connectance increased as the number of observed edges per species (sampling intensity) increased (Dormann et al., 2009). Consequently, Dormann et al. (2009) implemented null models in the bipartite R package to correct for sampling intensity. The BiMat MATLAB package uses a similar null model strategy to estimate statistical significance of network properties including nestedness, modularity and module structure (Flores et al., 2016). Connor, Barberán and Clauset (Connor et al., 2017) use Chung-Lu and Erdős-Rényi models to identify whether average path length, modularity, diameter and the clustering coefficient were significantly different for observed networks compared to networks generated with these models. In each of these examples, a network property of interest could be interpreted more meaningfully after comparison to random networks. In contrast to studying one network in particular, we adopted a null model strategy to specifically address the existence of conserved associations. While applications for network analysis such as NetConfer and setsApp also return intersections of networks (Morris et al., 2014; Nagpal et al., 2020), these applications do not assess whether this intersection is non-random.

In this manuscript, we introduce a null model strategy based on shuffled networks (Fig. 1). We illustrate the power of this strategy by identifying non-random CANs in sponge microbial networks. For these networks, we show that the CANs represent biologically relevant group-specific associations. We also demonstrate how degree-preserving randomized networks help identify betweenness centrality scores that result from the network’s connectivity pattern rather than from node degree distribution alone. These strategies have been implemented in a software toolbox that evaluates the significance of network or network property comparisons.

**Figure 1.**
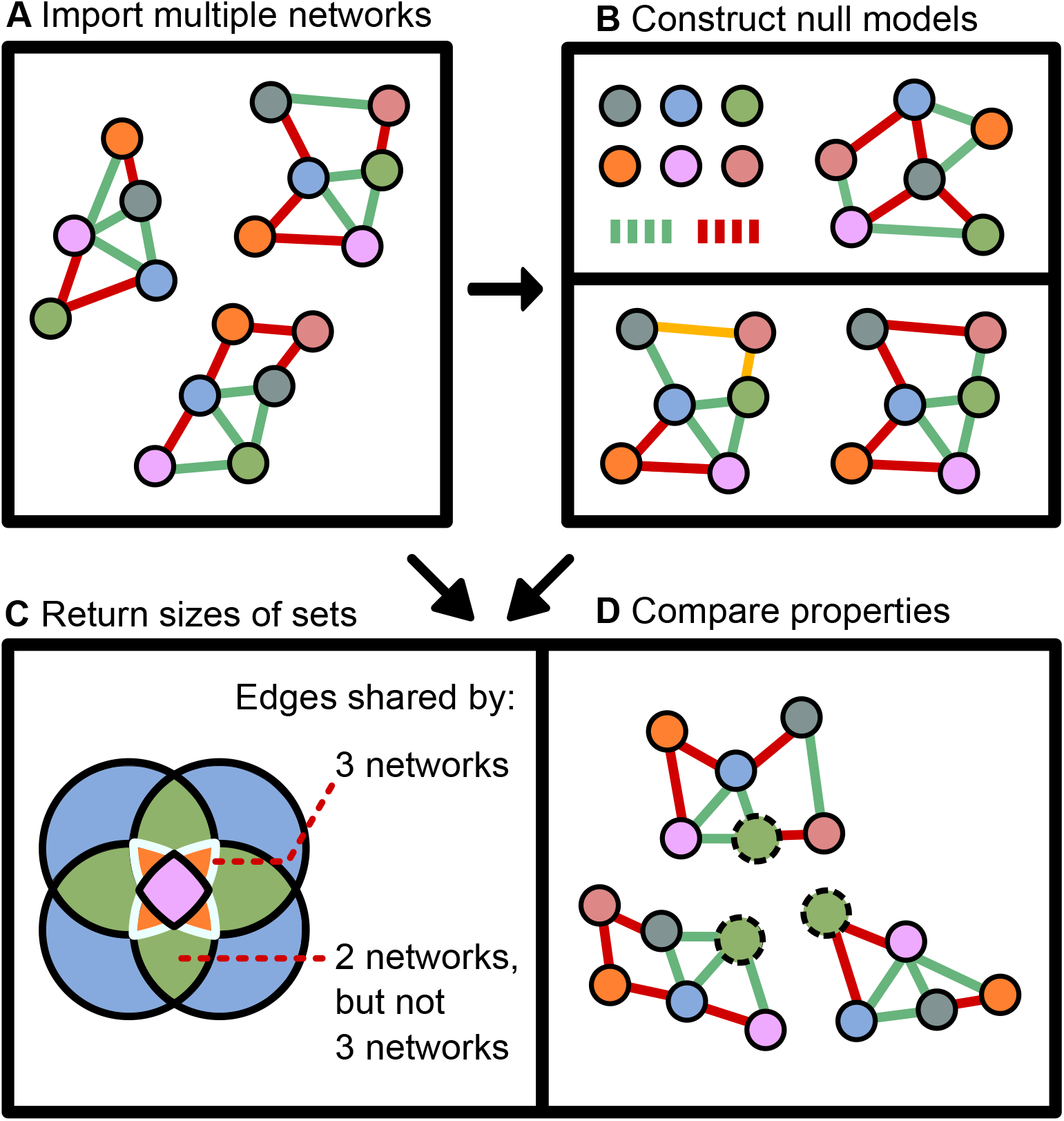
The anuran pipeline. **a** Networks are imported by the user. These networks can be ordered, and multiple groups of networks can be imported at the same time. **b** Random networks are constructed for each of the imported networks. These can be fully randomized, or they can preserve the degree distribution of the original network by swapping edges (highlighted in yello. They may also contain a synthetic core. **c** The toolbox returns sizes of sets (e.g. intersections of 3 networks) and sets of sets (e.g. the difference between the intersections of 2 networks and the intersections of 3 networks). These sets measure the overlap between specific numbers of networks. By comparing set sizes for observed networks and random networks, the significance of the set sizes can be assessed. **d** Network-level and node-level properties are computed for both the random networks and the observed networks, which allows assessing the significance of these properties within a group of networks and between multiple groups of networks.

## Methods

### Null model toolbox

We have developed a software toolbox, anuran, (**a** toolbox with **nu**ll models for identification of non-**r**andom patterns in **a**ssociation **n**etworks) that generates random networks and assesses properties of these networks. Three types of networks can be generated in the current implementation: completely randomized networks, degree-preserving networks and a variation of both networks that keeps a fraction of the edges fixed. For clarity, the completely randomized networks and degree-preserving networks without fixed edges are referred to as negative control networks, and the same networks with fixed edges are referred to as positive control networks in the remainder of the manuscript. For the completely randomized model, a network is initialized with the same nodes as the input network. Edges are then added randomly until the total edge number is equal to the number of edges in the input network. For the degree-preserving model, edges are swapped rather than removed and added back to the network, so that two edges (a,b) and (c,d) become the new edges (a,c) and (b,d). Hence, the model preserves the degree distribution found in the input network and each node has the same degree as it has in the original network, but other centralities such as the betweenness centrality can change. The user specifies both the number of random networks generated for each network (by default 10) and the number of sets of these networks (by default 50) that are sampled to calculate set sizes.

The toolbox can generate random networks with a fraction of fixed edges, which serve as a type of positive control. First, a fraction of edges is extracted from the total union of edges across all networks. For fully randomized networks, these edges are first added, then edges are added until the total number of edges in the original network is reached. For the degree-preserving randomized networks, negative control networks (with preserved degree) are first generated. Then, for each edge in the fixed core, the algorithm attempts to find two edges that can be swapped so the fixed edge is created. If this fails, a random edge is deleted and the fixed edge is introduced, so the degree is not exactly preserved. To swap the edges successfully, it is necessary that each of the nodes participating in a fixed edge has another edge not part of the fixed core. As a result, the degree distribution can change significantly for networks where nodes in the fixed core are disconnected, or where the fixed core is very large compared to the positive control network. However, for all positive control networks, edges in the core are always added.

It is possible to include nodes without significant associations in the network file as disconnected nodes (orphan nodes). This can be done by simply supplying the network file with the orphan nodes included as nodes without any edges. In this case, the random model more accurately reflects a situation where associations are randomly selected from the number of taxa in the ecosystem. However, the degree-preserving networks will not change, since an orphan node has a degree of 0 and will retain a degree of 0 in the null model. The inclusion of orphan nodes leads to different estimates for set sizes for the random model that may lead to an overestimation of the significance of a CAN. We expect that many nodes present in the abundance file will not be prevalent enough, not abundant enough or not variable enough to ever have a significant association in the first place. Therefore, we ignored the presence of disconnected nodes in our case study.

The toolbox has been implemented in Python 3.6 and consists of both an application programming interface (API) and command line interface (CLI) that carries out a complete analysis. Functions from the API can be used to integrate the toolbox into other projects and pipelines and have been structured in a modular manner so customization of the toolbox is straightforward. For example, users can incorporate any centrality or network property implemented in another library, e.g. NetworkX, without making large changes to the toolbox (Hagberg et al., 2008). Currently, the CLI pipeline assesses set sizes, (rank-transformed) betweenness, degree and closeness centrality scores and several network-level properties: degree assortativity, connectivity, diameter, radius and average shortest path length (Fig. 1). NetworkX implementations of these centrality calculations were used (Hagberg et al., 2008).

The software uses a set-of-sets approach to identify CANs. A set is a specific collection of edges, such as the intersection set, which is the collection of edges present across multiple networks. The CANs are identified as differences of specific intersection sets. Hence, the toolbox specifically identifies sets and sets-of-sets that are likely to be of interest for microbial association networks. These sets represent collections of edges that are only present in one specific fraction of networks and distinguish between less-conserved and more-conserved edges.

An example with 4 networks is illustrated with a Venn diagram (Fig. 1c). The intersection containing all edges in at least 2 networks is the green part of the Venn diagram plus the other colours inside this part. To obtain the difference of the intersections, the set that includes one or more additional networks is subtracted from the intersection set that includes fewer networks. These sets are referred to as combinations of intersections with fractions or integers. Therefore, the intersection 0.5 refers to all intersections of 50% of the networks, while the intersection 3 refers to intersections that contain edges present in at least 3 networks. Similarly, set-of-sets are identified by a combination of intersection numbers: the set-of-sets 6 → 10 refers to the difference of intersection 6 and intersection 10 and therefore contains no edges present in at least 10 networks. For most analyses, the difference of intersections is preferred over intersections since the intersections are nested. By taking the difference, it is possible to distinguish between more and less conserved associations.

The equations for differences and *k* -intersections for groups of *n* networks are given below. The equations only refer to edge sets *E*, so they do not apply to numbers of matching nodes. The difference is the union of all sets *D*_*i*_ for 1 up to *n* networks, where the sets *D*_*i*_ contain all edges *x* present in an edge set *E*_*i*_ but not in the union of all other edge sets:

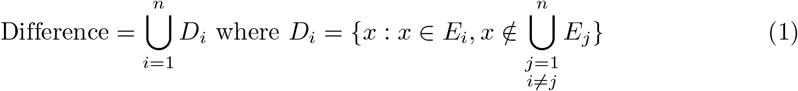

The *k* -intersections are unions of intersections *S*_*I*_. These intersections *S*_*I*_ are sets of groups of edge sets, where the groups *I* are *k* -permutations of *n* and *E*_*i*_ is a single edge set in *I*. Hence, for a total number of edge sets *n*, each of the groups *I* have size *k* and the collection of all possible groups is indicated as 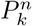. For the 4-intersection for a group of 40 edge sets, the size of 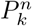 can be calculated as the binomial coefficient 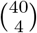 This mathematical representation is not implemented directly in the software, as the software simply takes the set of all edges present in at least 4 networks and therefore ignores network identity.

Hence, a *k* -intersection is the union of all intersections *S*_*I*_ for *I* in 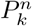:

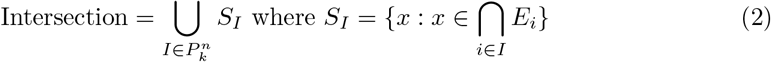

Since edges present in at least *k* networks but not in *m* networks represent less-conserved edges, the difference of the intersections is calculated to distinguish between less-conserved and more-conserved edges. The difference of two intersections *k* and *m*, with *S*_*I*_ and *S*_*J*_ defined identically to *S*_*I*_ in the equation above, is then given below:

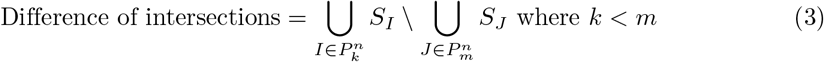

To compare observed set sizes to set sizes of random networks, the Z-score test is carried out, which identifies set sizes in the input networks that are outside the range of set sizes inferred from groups of random networks. The scipy normaltest implementation (Virtanen et al., 2020) of D’Agostino’s and Pearson’s omnibus normality test was used to test for both kurtosis and skewness (d’Agostino, 1971; Pearson et al., 1977). Since this test requires at least 20 observations, a warning is issued if the number of random networks needs to be increased.

The toolbox can also calculate centrality scores for groups of networks. To ensure that centralities are not biased by edge number, these are first converted to ranks before a Mann-Whitney *U* test is used to assess whether the distributions of ranks are similar across groups of observed networks and random networks. The comparisons to random networks are repeated a number of times and parameter-free p-values across all comparisons are calculated from the number of successful Mann-Whitney *U* tests. By default, multiple-testing corrections are carried out on these p-values to correct for the number of taxa. Any of the corrections available in the multipletests function from the Python package statsmodels are available (Seabold & Perktold, 2010). The approach for network-level properties is similar, with the software currently supporting assortativity, connectivity, diameter, radius and the average shortest path length. If the networks are ordered, the toolbox can calculate Spearman correlations of these properties to the network order. For example, users could supply networks constructed across a pH gradient and the software would test whether there is a correlation between the network indices (from 1 to the network number) and centralities or other network properties. The results of these analyses are exported to tab-delimited files so they can be further analyzed and visualized in the user’s preferred statistical environment.

Finally, the toolbox includes an option for resampling networks. This option samples a subset of networks for the specified subsets sizes. In this way, the resulting data shows how trends in set sizes change as the number of networks is increased. The resulting data can be interpreted as a form of rarefaction curve, where a flattening of the curve suggests that sufficient networks have been collected to identify all edges present in a specific fraction of networks.

### Case study

QIIME-processed data was downloaded from Moitinho et al. (Moitinho-Silva, Nielsen, et al., 2017). Samples with fewer than 1000 counts were removed and the samples were rarefied to even depth. After rarefaction, the abundance data were first filtered for 20% taxon prevalence across all samples, then once more to ensure 20% prevalence across the different orders. Counts for removed taxa were retained to preserve the sample sums. After excluding host orders with fewer than 50 samples, 10 orders remained. CoNet was then used to infer association networks (Faust & Raes, 2016). Edges were generated with Pearson correlation, Spearman correlation, mutual information, Bray-Curtis dissimilarity and Kullback-Leibler distance. Edges were included if at least one method reached significance; only edges with a combined Q-value below 0.05 (estimated using a combination of permutation and bootstrapping) were retained. The CoNet CANs were inferred with anuran generating 20 negative control random networks per host order and resampling these 100 times. For the positive controls, 20 network groups were generated with a core size equal to 20% of the union of edges at 20% prevalence (edges present in at least 2 networks) and at 50% prevalence (edges present in at least 5 networks). Set sizes and centralities with a p-value below 0.05 for comparisons to values from random networks were considered significantly different from the random networks. CoNet networks were compared to FlashWeave networks (Tackmann et al., 2019). FlashWeave was run as FlashWeave-S (sensitive set to true and heterogeneous to false), with all other settings set to the default. To compare FlashWeave networks to CoNet networks, *anuran* generated five randomized networks per order-specific network and resampled these five times.

Prior research indicated that microbial abundance was a significant driver of community structure (Moitinho-Silva, Steinert, et al., 2017). Therefore, taxa in the CAN were compared to taxa reported as indicators of high microbial abundance or low microbial abundance (Moitinho-Silva, Steinert, et al., 2017). CAN network clusters were identified with manta (Röttjers & Faust, 2020b), as this algorithm has been designed to handle negative edges in the CAN. To run the clustering algorithm, default settings were used, except the number of iterations and permutations, which was set to 200. A Chi-squared test was used to compare HMA-LMA predictions to CAN cluster assignments (*α*=0.05).

## Results

We analyzed 10 sponge order-specific networks that we inferred from Sponge Microbiome project data (Moitinho-Silva, Nielsen, et al., 2017). Due to their sessile lifestyle, sponges protect themselves from overgrowth, predation and competition through production of bioactive compounds (Proksch et al., 2010). Such compounds may be produced by the sponges themselves or by their microbial symbionts (Wijffels, 2008). Consequently, sponges may be expected to harbour symbiotic species that improve sponge health. While their open connection with their surroundings suggests that part of their microbiome may be transient, stable core microbiomes have been identified (Thomas et al., 2016). Therefore, our toolbox provides an opportunity to investigate conserved associations across sponges.

Networks were constructed with CoNet (Faust & Raes, 2016). These networks had a median edge number of 137, with the smallest network containing 56 edges and the largest 1735 edges. We confirmed that a different network inference tool, FlashWeave, was able to recover many of the same associations despite large differences in network size (Fig. 2)(Tackmann et al., 2019). Likely, the difference in edge number can be attributed to the tests for conditional independence used by FlashWeave to reduce the number of indirect edges.

**Figure 2.**
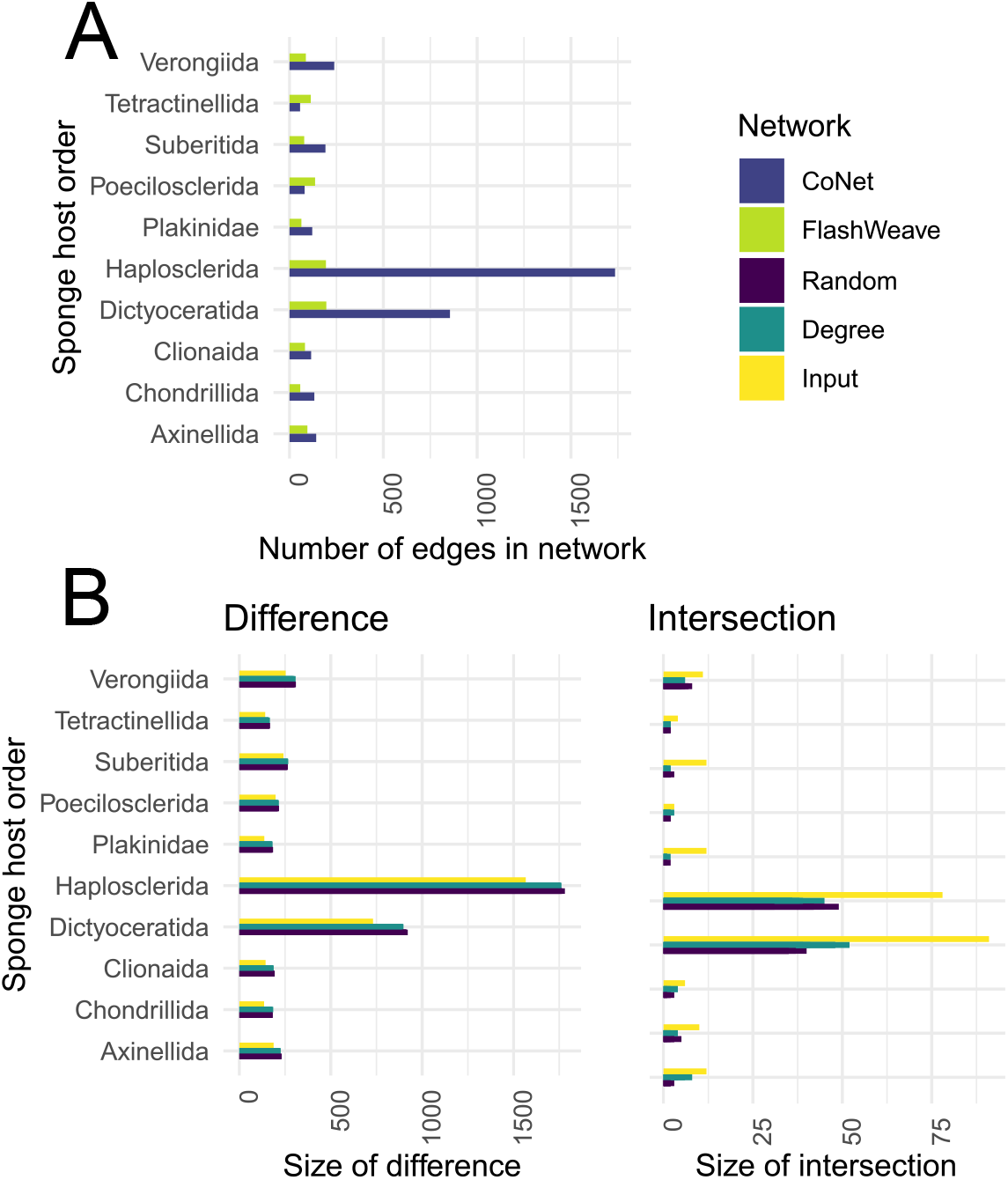
Comparison of FlashWeave and CoNet networks on sponge order-specific networks. Host order-specific networks were generated with both CoNet and Flashweave from samples collected for the Sponge Microbiome Project. The labels on each figure refer to the sponge order. **a** Sizes of the FlashWeave and CoNet networks. **b** Difference and intersection of the FlashWeave and CoNet networks. The different bars represent the observed set size for the observed data (Input), the fully randomized networks (Random) and the degree-preserving randomized networks (Degree).

**Figure 3.**
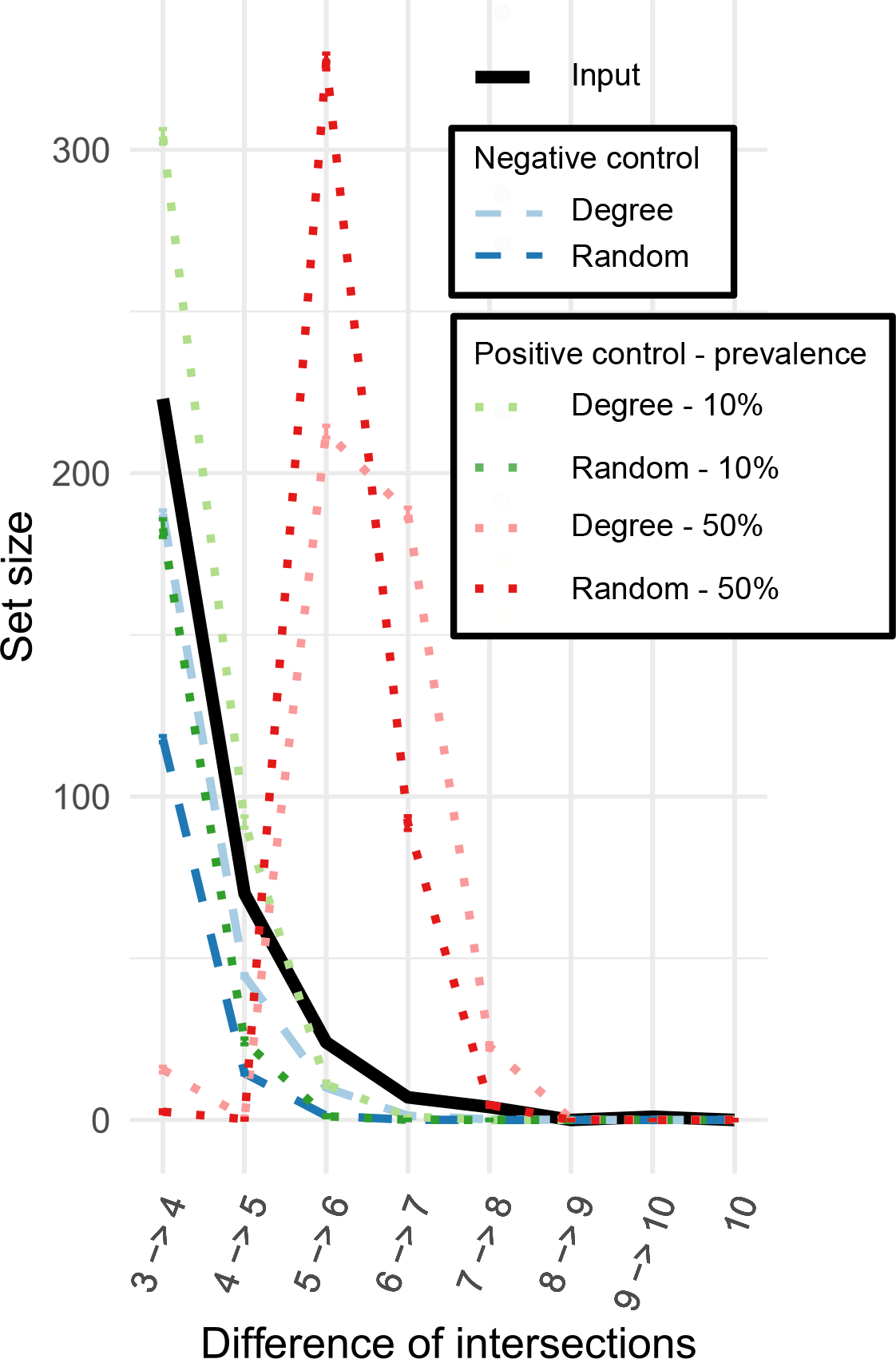
Set sizes across networks for 10 sponge networks and randomizations of these networks. The set size is the number of edges present in a particular number of networks. The set size is shown for a set-of-sets of network intersections, meaning that the set-of-sets 4 → 6 is calculated as the number of edges in 4 or 5 networks with all edges in at least 6 networks removed. Each network was generated for a different host sponge order for which at least 50 samples were available. These networks were then randomized either with the same degree distribution (Degree) or without this distribution (Random). Moreover, each of the networks was randomized with preservation of a part of the input network for a subset of the randomizations as a positive control. Hence, the positive control Degree networks are randomized versions of each input network with 20% of the union of associations present in at least 20% of observed networks or at least 50% of observed networks. Error bars represent the standard error across different combinations of random networks. For sets of edges present in up to 6 networks, the set size of the input networks deviates significantly from the set size from those of random networks with or without degree preservation.

Intersection differences up to 6 networks were significantly larger than differences generated from the randomized and degree-preserving negative control networks (p*<*0.0001) (Fig. 3). However, the positive controls with a core conserved across 50% of networks had a much larger set size at 5 networks. Therefore, the CAN was constructed from all associations present in 3 out of 10 networks (Fig. 3). Prior work suggests that most of the variation in a bipartite sponge-bacteria network could be attributed to differences between bacterial abundance: high microbial abundance (HMA) versus low microbial abundance (LMA) (Moitinho-Silva, Nielsen, et al., 2017). Supplementary data from Moitonho-Silva et al. (Moitinho-Silva, Steinert, et al., 2017) were used to identify taxa in network clusters that were significantly more or less abundant in HMA compared to LMA sponges in their work. Indeed, we found that HMA and LMA assignments were different across the three clusters (Chi-squared test, p=0.006), with cluster 0 containing more HMA-associated phyla and cluster 1 containing only LMA-associated phyla. This suggests that the CAN contains several phyla that have previously been identified as indicators of HMA-LMA status.

**Figure 4.**
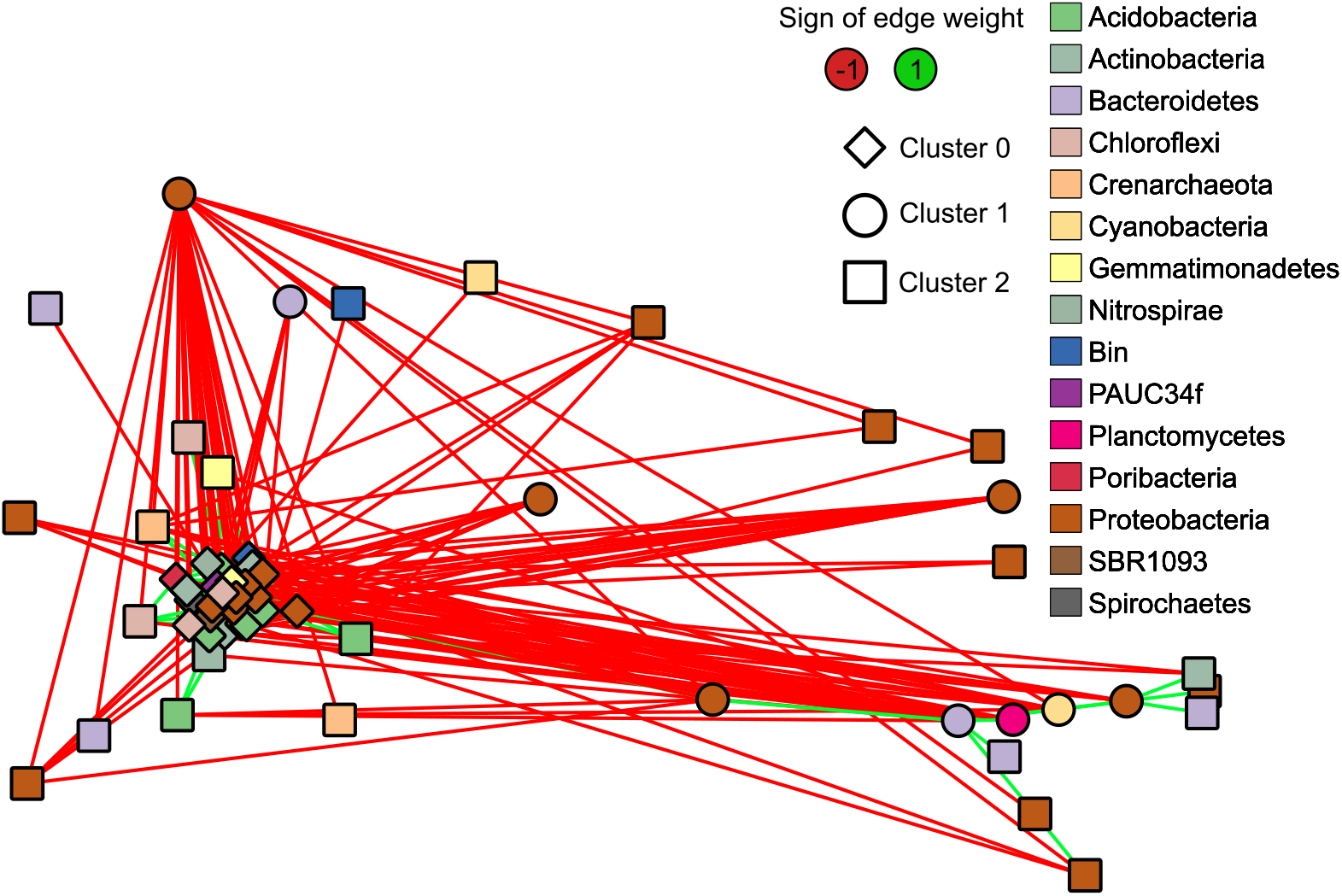
Core association network (CAN) constructed from associations present in at least 3 orders of sponges. Networks were generated with CoNet from samples collected for the Sponge Microbiome Project (Moitinho-Silva, Nielsen, et al., 2017) and were constructed using samples from a single order of sponges. Associations present in at least 3 networks were included in the CAN. Node colour was mapped to phylum and the manta clustering algorithm was used to identify clusters in the network.

In addition to a CAN, we investigated whether the networks contained taxa with consistently lower or higher centrality scores compared to the randomized networks (Additional File 1: Figure S7). Since no taxa had significantly different scores, we instead investigated three taxa with uncorrected p-values below 0.3. All three taxa had lower degree centralities compared to the randomized networks: these were a taxon assigned to the phylum Acidobacteria (p=0.188), a taxon belonging to the SAR202 clade (p=0.119) and another assigned to the Alphaproteobacteria (p=0.129). All p-values were 0.99 after Benjamini-Hochberg multiple-testing correction for taxon number. Overall, centrality scores were too variable across networks to conclude whether they were significantly different.

## Discussion

Researchers can use properties of microbial association networks to describe trends in microbial communities. Such properties can include modularity, the existence of a CAN, a high degree (hub nodes) or any other property that can be calculated from the network. Frequently, these are considered to mirror how the studied community is structured. In ecological networks, nestedness may even predict ecosystem robustness (Burgos et al., 2007). Hence, robust estimates of these properties can yield valuable information on community structure.

We chose to use two types of network null models that represent two extremes in terms of constraints. The randomized null model is not constrained in terms of network structure (apart from node and edge number), while the degree-preserving models may be overly constrained especially if the degree centrality is a meaningful representation of a biologically relevant property, such as a taxon’s generalist lifestyle. Moreover, these models make no assumptions on the nature of associations. Therefore, it is unknown whether an association in a CAN reflects a biotic interaction, as these associations can also result from similar taxon responses to the environment or to other organisms. The effects of biotic interactions can be better studied through other methods, for instance joint species distribution models (Ovaskainen et al., 2017; Tikhonov et al., 2017).

Additional assumptions could be included to further improve the ability of null models to identify striking patterns. However, more complex null models are not well-established when it comes to the analysis of microbial association networks. Candidates include the Barabási-Albert (Albert & Barabási, 2002) and Klemm-Eguíluz network models (Klemm & Eguiluz, 2002), which describe mechanisms of network growth and could therefore identify networks not generated in accordance with such mechanisms. Yet, a network model that assumes a particular growth mechanism may not be appropriate for association networks. Few associations in an association network are expected to represent interactions (Freilich et al., 2018). As a result, mechanisms for network growth may apply to the underlying interaction networks, but not directly to association networks. If a null model includes an assumption not known to be true, the null model becomes a pseudo-null model (Bausman & Halina, 2018). This could wrongly lead researchers to conclude that there is no relevant biological effect in addition to the effect described in the pseudo-null model. A comparison to these network models therefore addresses whether properties are significantly different compared to networks generated according to specific rules of network growth, but it cannot address the non-randomness of network properties. Since this toolbox has been developed to find non-random trends in association networks, we chose network null models that make no assumptions on network growth (e.g. preferential attachment) because 1) we cannot be sure that such assumptions hold for interaction networks and 2) these null models do not take additional processes into account (such as environmental influence) that likely shape association networks.

As one of the null models in anuran preserves the degree distribution, comparisons to networks generated from those null models support statements on correlations between degree and other centralities. While we did not fully explore the effects of other properties, such as the fraction of realized edges or network topology, these topics have been discussed previously in methodological studies on ecological and social networks (Dormann et al., 2009; Faust, 1997; Valente et al., 2008; Valente & Foreman, 1998). However, we do want to emphasise that they deserve similar attention in the context of microbial association networks. For example, Agler et al. (2016) defined hub taxa as those taxa that had both higher closeness, betweenness and degree centrality than other taxa (Agler et al., 2016), but such measures may be strongly correlated in dissortative networks, where nodes with high degree are more likely to connect to nodes with low degree (Goh et al., 2003). Our analysis of centrality rankings further supports the observation that betweenness and degree centrality can be correlated, as we found that taxa did not have betweenness centrality rankings significantly different from betweenness centrality rankings observed for the degree-preserving random networks.

On networks constructed from sponge order-specific taxon abundances, the set-of-sets approach identified a large CAN. This CAN was significantly different from the CAN observed for random networks. We suspected based on prior work that this large CAN could arise from a major partition in sponge symbiotic relationships: sponges tend to either have high or low microbial abundance (Gloeckner et al., 2014). Several taxa have been identified as indicators of this divide and many of those indicators were also found in the CAN (Moitinho-Silva, Nielsen, et al., 2017). Although HMA-LMA status is not strictly phylogenetically conserved across most sponges (Gloeckner et al., 2014), associations between taxa that relate to this status appear to be conserved across at least a subset of sponge orders (Figure 5). Hence, the set-of-sets analysis suggests that some of the dynamics responsible for the HMA-LMA discrepancy are shared across different orders of sponges. Therefore, the flexible CAN threshold that we adopted has the potential to identify group-specific dynamics that can reveal more about community structure.

**Figure 5.**
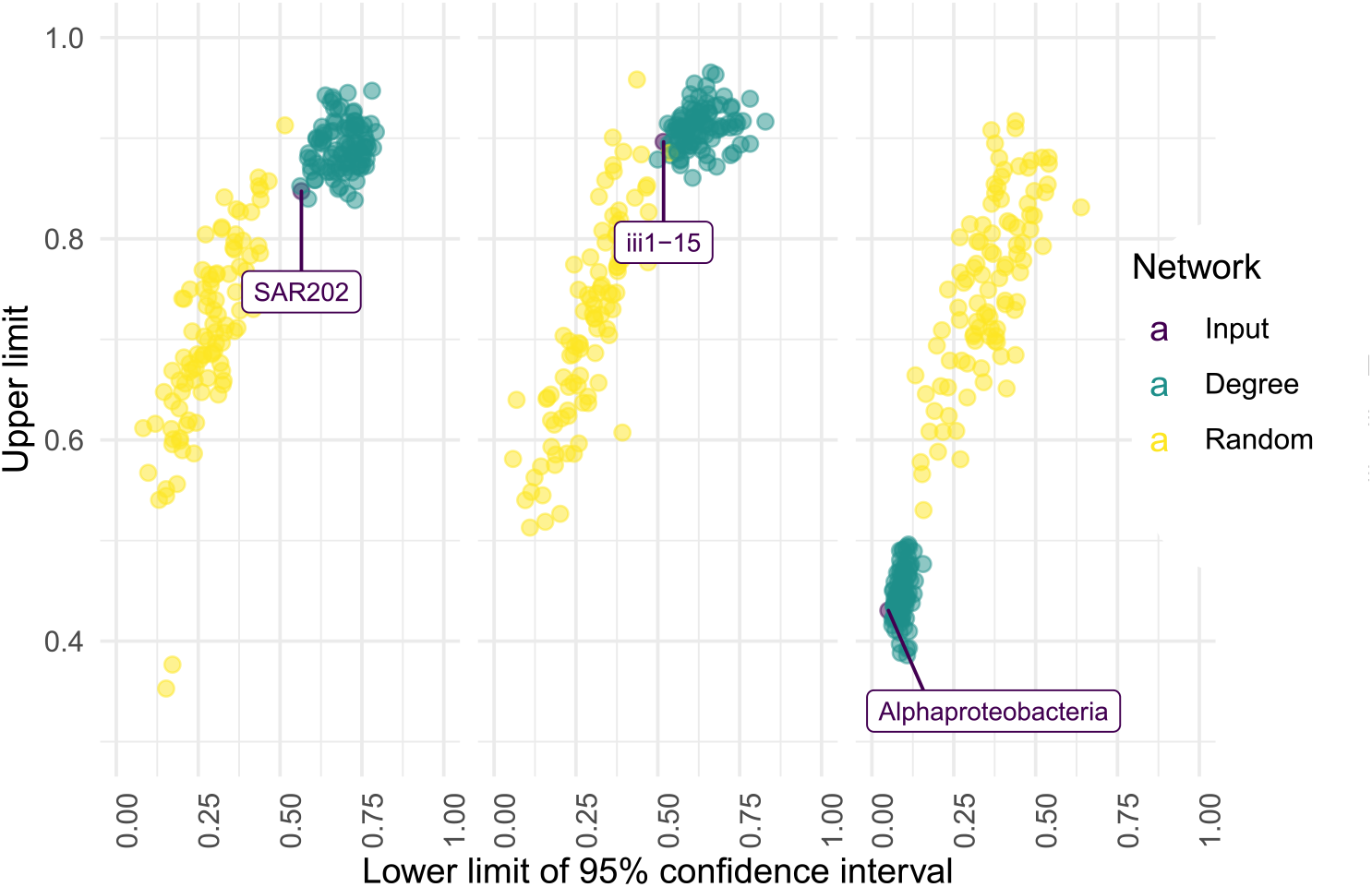
Confidence intervals of betweenness centrality rankings for three taxa found in sponge order-specific networks. Estimated upper limit and lower limits for the 95% confidence intervals of betweenness centrality computed for each taxon across a group of networks and null models generated from these networks. Taxa that consistently have high centralities have narrower confidence intervals and are therefore located in the upper right of the plot, while taxa with consistent low centralities are located in the bottom right. Null models either had the same degree distribution (Degree) or were fully randomized (Random). The shown taxa were selected because they had a p-value below 0.3 for any of the permutation tests comparing closeness, betweenness or degree centrality rankings. The taxa labelled SAR202, iii1-15 (a clade of Acidobacteria) and Alphaproteobacteria had p-values of 0.119, 0.188 and 0.129 for the degree centrality tests compared to the fully randomized models.

## Conclusions

Microbial networks have become a popular method for the analysis of microbiome data despite their low accuracy. With anuran, we have introduced a tool that can aid in the comparison of multiple noisy networks through analysis of random networks. These networks permit users to compare centrality rankings, to infer core sizes and to construct correlations across ordered networks. Our set-of-sets approach uses null models to find conserved patterns across groups of networks. We demonstrate that such a set-of-sets approach, by using an alternative null model strategy, can also detect redundancy between network properties such as degree and betweenness centrality. Therefore, anuran is one of the first dedicated tools for meta-analysis of noisy networks with null models. We expect this null model suite to be a valuable benchmarking tool in the analysis of microbial and other networks.

## Data availability

All scripts and software, including scripts to generate the figures in this manuscript, have been deposited to Zenodo at https://zenodo.org/record/4030380 (Röttjers et al., 2020). An up-to-date version of the software is being maintained on a Github repository: https://github.com/ramellose/anuran (Röttjers & Faust, 2020a). All materials are available under the Apache 2.0 license.

## Funding

This project was supported by the KU Leuven under grant number STG/16/006. KF has received funding from the European Research Council (ERC) under the European Union’s Horizon 2020 research and innovation programme under grant agreement No 801747. The Raes laboratory is supported by the VIB Grand Challenges programme, KU Leuven, the Rega Institute for Medical Research, and the FWO EOS program (30770923). DV is supported by a post-doctoral fellowship from the Research Foundation Flanders (FWO Vlaanderen).

## Notes

https://zenodo.org/record/4030380#.X3rQPedx1hE

### Competing Interest Statement

The authors have declared no competing interest.

